# Integrating ecological monitoring and local ecological knowledge to evaluate conservation outcomes

**DOI:** 10.1101/2021.05.31.446466

**Authors:** Michelle María Early-Capistrán, Elena Solana-Arellano, F. Alberto Abreu-Grobois, Gerardo Garibay-Melo, Jeffrey A. Seminoff, Andrea Sáenz-Arroyo, Nemer E. Narchi

## Abstract

Successful conservation of long-lived species requires reliable understanding of long-term trends and historical baselines. Using a green turtle (*Chelonia mydas*) foraging aggregation in the northern Gulf of California, Mexico as case study, we integrated scientific monitoring data with historic catch rate reconstructions derived from Local Ecological Knowledge (LEK). Models fit to LEK and monitoring data indicate that turtle abundance is increasing, but only after ~40 years of safeguarding the species’ nesting and foraging habitats in Mexico. However, as population declines occurred 75% faster than increases, and current abundance is at ~60% of historical baseline levels, indicating the need for sustained, long-term conservation actions. This study demonstrates the potential of linking LEK and ecological science to provide critical information for conservation, by establishing reference baselines and gauging population status, while promoting equitable and sustainable futures for local communities.

## Introduction

Successful conservation of highly migratory, long-lived species such as sea turtles requires efforts and policies implemented across large spatio-temporal scales, along with locally-grounded data and practices (Mazaris et al. 2017; Vierros et al. 2020). Local Ecological Knowledge (LEK) can contribute to sound conservation and management practices by establishing local baselines and recovery targets; evaluating population status; integrating the cultural dimensions of human-environment interactions; and supporting equitable and inclusive practices (Early-Capistrán et al. 2020a; Lee et al. 2018; Poe et al. 2014). LEK-based approaches are particularly important for taxa impacted by small-scale fisheries or subsistence hunting, for which technical or baseline abundance data may be scarce or un available (cf. Sáenz-Arroyo & Revollo-Fernández 2016; Selgrath et al. 2018).

East Pacific green turtles (henceforth, green turtles) are a regionally-distinct population segment of the cosmopolitan green turtle (Seminoff et al. 2015). These marine megaherbivores are ecosystem modifiers, with vital roles in marine food-web and ecosystem dynamics (cf. Christianen et al. 2021). They are highly migratory and occupy a broad range of habitats across different life stages, primarily tropical reproductive areas and tropical or warm-temperate foraging areas separated by hundreds or thousands of kilometers (cf. Seminoff et al. 2015). Globally, monitoring is heavily skewed towards nesting habitats where only breeding females and egg output are quantified (cf. Seminoff & Shanker 2008). However, coastal and neritic foraging habitats incorporate a broad range of turtle age classes and both sexes, providing insights into population dynamics and trends crucial for informing conservation policy (Bjorndal et al. 2005; Seminoff & Shanker 2008; Seminoff et al. 2003).

Like other sea turtle species, green turtles are highly susceptible to over-exploitation due to their complex life-history and prolonged maturation time. Worldwide, the long history of commercial exploitation of meat and eggs has depleted or extirpated several green turtle stocks (cf. Chaloupka et al. 2008). The Baja California peninsula provides a unique case study with a robust historical baseline within living memory (the 1950s) (Early-Capistrán et al. 2018). Green turtles are a cultural keystone species (Garibaldi & Turner 2004) used over millennia by the region’s inhabitants, with fundamental roles as food and medicine. Small human populations and geographic isolation kept captures sustainable until the mid-twentieth century. From 1960-1980, intensive commercial fishing supplied green turtle meat to fast-growing cities along the U.S.-Mexico border, driving foraging populations to near extinction (Early-Capistrán et al. 2018). This process was coupled with intense egg collection in the populations’ tropical reproductive habitats (Delgado-Trejo & Alvarado Díaz 2012; Seminoff et al. 2015) (Figure 1). Long-term conservation and research began in the late 1970s as green turtles diminished, and in the early 1980s the fishery collapsed (Early-Capistrán et al. 2020a; Seminoff et al. 2008) (Figure 2).

**Figure 1:**
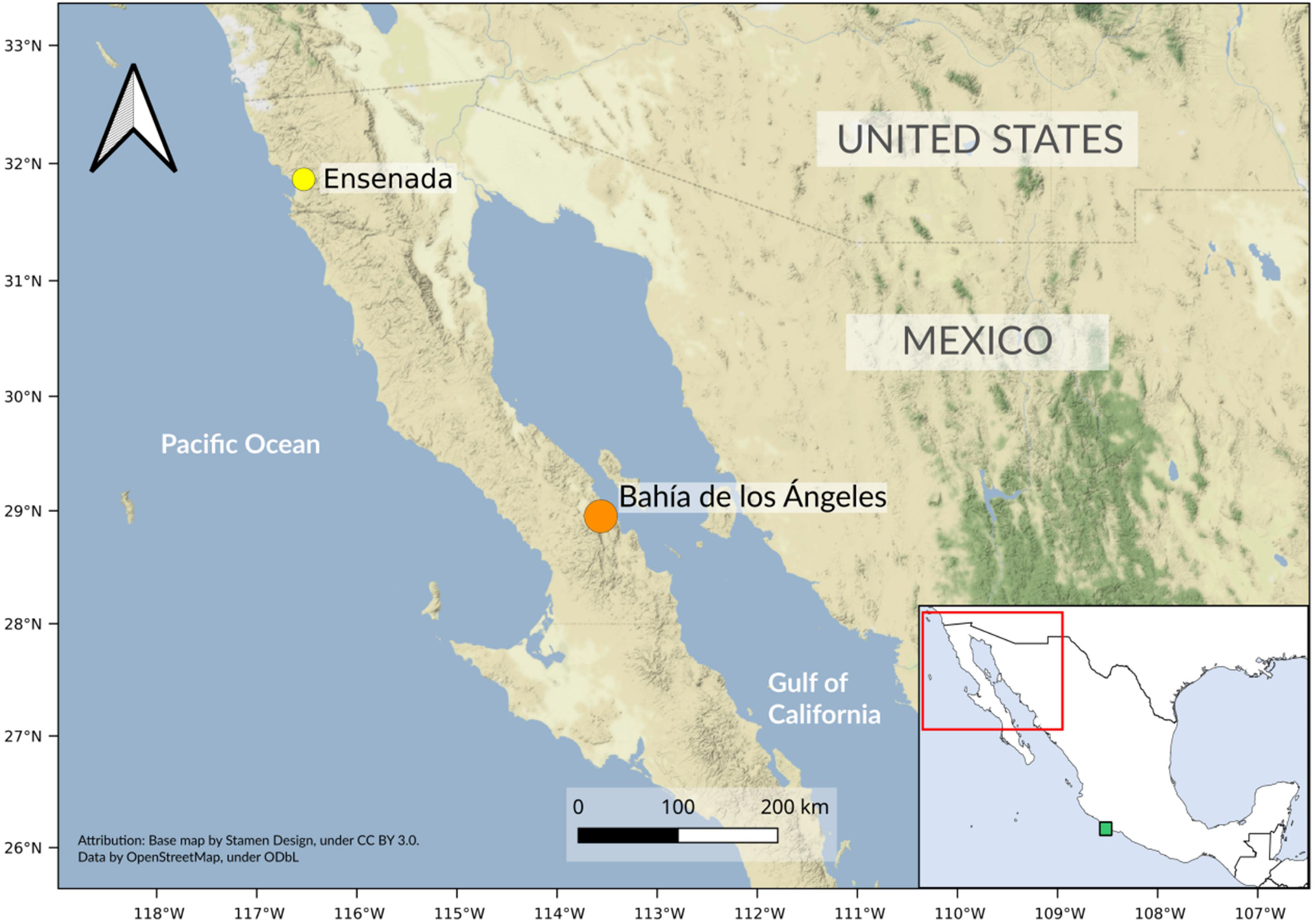
Map showing the location of the study site at Bahía de los Ángeles, Baja California, Mexico (orange circle) and Ensenada, Mexico (yellow circle), the primary market for green turtle catches during the historical commercial fishery. The green square in the inset map shows the index nesting site in Colola, Michoacán. At the time of this study, the village of Bahía de los Ángeles had a population of ~500 people, with economic activity centered on small-scale fishing and seasonal tourism. Map produced in Qgis using Stamen Terrain Basemap (CC BY 3.0) and data from OpenStreetMap (ODbL).

**Figure 2:**
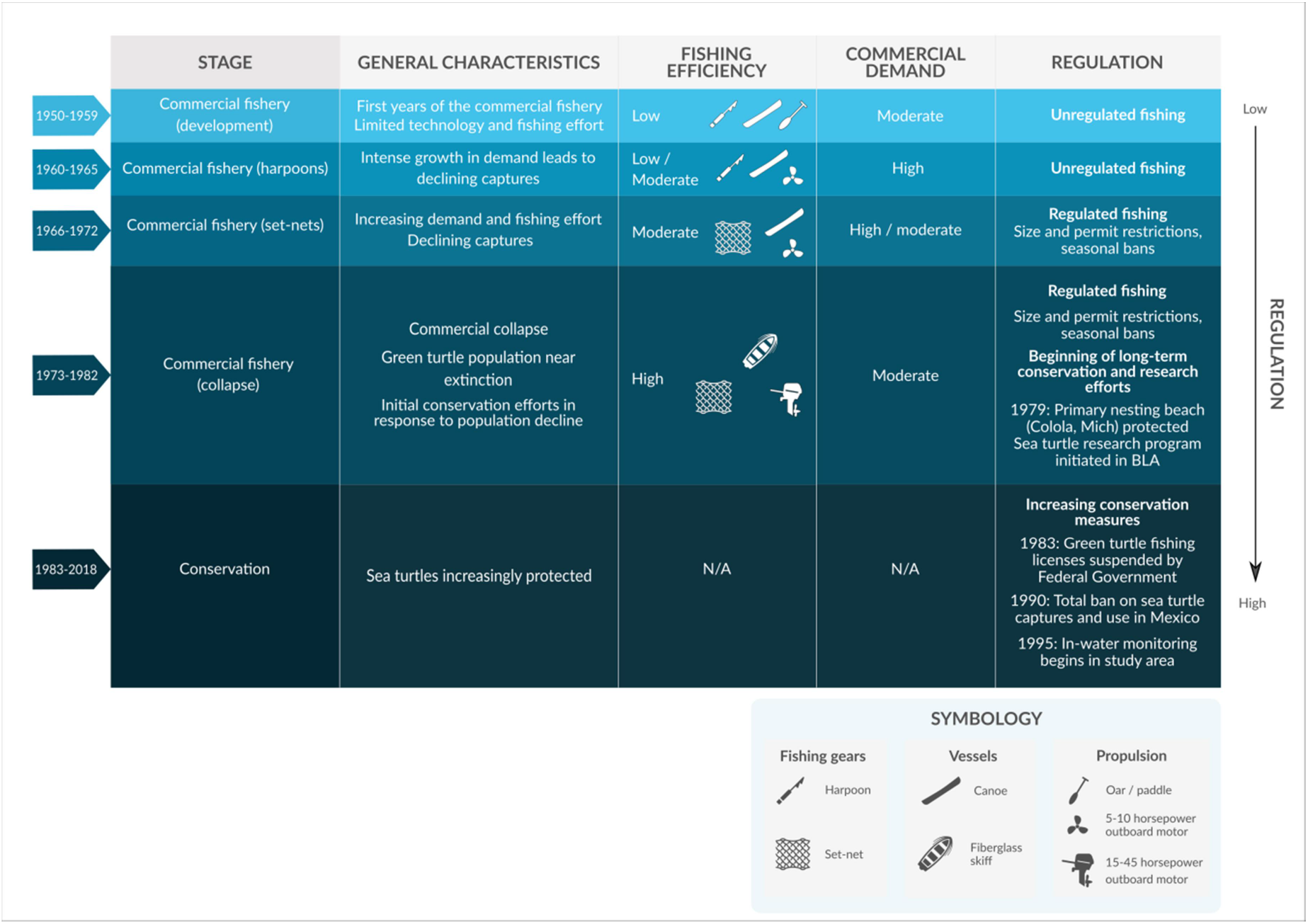
General chronology of changes in the green turtle fishing and conservation in Bahía de los Ángeles. Fishing efficiency increased along with demand between 1960 and 1980, leading to population collapse. Long-term conservation and research began in the late 1970s in response to diminishing populations. Fishing permits for green turtles were suspended by the Federal Government in 1983 as the fishery collapsed. Regulations and conservation measures across green turtle habitats increased from the late 1970s onward, reaching full legal protection after the total ban in 1990 (Early-Capistrán et al. 2020; Márquez 1996; Seminoff et al. 2008).

Local, regional, and international conservation efforts, including nesting beach and habitat protection, bans or restrictions on direct use, and by-catch regulations, have successfully reversed declines in some populations, including green turtles in the east Pacific (Broderick et al. 2006; Mazaris et al. 2017). Green turtles are classified as endangered under Mexican law and by the IUCN, and all sea turtle exploitation in Mexico has been banned since 1990 (Diario Oficial de la Federación 1990; IUCN 2019). Decades of nesting beach and habitat protection across the species’ range in Mexico have led to population increases since the early 2000s (cf. Seminoff et al. 2015).

We integrated scientific monitoring data with historical catch rate reconstructions generated collaboratively with local fishers by documenting and quantifying LEK, resulting in (to our knowledge) the longest available standardized time-series (1952-2018) worldwide for a sea turtle foraging habitat. We aim to understand long-term change, contextualize current local abundance compared to historical baselines, and help establish future directions for conservation and management policies. Our innovative, long-term approach can help link ecological science and LEK, promoting conservation processes that harness the collective capacity of local and scientific communities (Game et al. 2015).

## Methods

### 2.1 Study site

Bahía de los Ángeles (BLA), Baja California, Mexico (28.951917°, −113.562433°) is a warm-temperate green turtle foraging area in the Gulf of California (Figure 1). This index site hosts significant in-water aggregations of turtles and systematic long-term scientific monitoring (Early-Capistrán et al. 2020a), initiated in 1979 through government-sponsored efforts (Seminoff et al. 2008). In-water monitoring began in 1995 and has continued as a joint effort between government, academic, and non-governmental institutions (Figure 2). Green turtles do not nest in BLA. The index nesting beach for this population is in Colola, Michoacán, Mexico, ~1500 km southeast of BLA (18.297392°, −103.410956°) (Seminoff et al., 2003). Establishing direct connectivity with stock-specific nesting rookeries is challenging, as green turtle foraging areas incorporate individuals from multiple genetic stocks (Dutton et al. 2019; Seminoff et al. 2015). However, Colola is the only green turtle nesting site in the Northeast Pacific with long-term data (>30 years), and accounts for 56-71% of green turtles in the Gulf of California’s foraging areas (Delgado-Trejo 2016; Koch 2013).

### 2.2 LEK-derived data (1952–1982): commercial fishing

We used an existing dataset of LEK-derived catch-per-unit-effort (CPUE) estimates to establish baseline abundance and analyze long-term change before scientific monitoring (Early-Capistrán et al. 2020b) (Figure 2). We compiled data in an on-going transdisciplinary research process with the BLA community initiated in 2012. We documented, corroborated, and classified LEK through ethnography; synthesized through coding, indexing, and heuristics; and integrated feedback processes with statistical analyses. We standardized all CPUE estimates to one 12-h in-water set for a single 100m net. Standardization (i) removed the majority of variation not attributable to abundance changes (e.g., fishing gear types, fleet characteristics, spatial dynamics); and (ii) generated estimates compatible with monitoring data. To evaluate central tendencies, we used annual mean CPUE for all analyses (Supporting Information: 1, Figure S1, Table S1). LEK data were shown to be statistically reliable and generated robust models of population change (Early-Capistrán et al. 2020a). See Early-Capistrán and collaborators (2020a) for detailed methodological processes.

### 2.3 Scientific monitoring (1995-2018): conservation

In-water monitoring began in 1995 after population collapse. Using CPUE as an index, turtles were captured with set-nets of the same design as those used in commercial green turtle fishing (Seminoff et al. 2003) (Supporting Information: 2). Monitoring data were provided by author J.A. Seminoff (2003; NOAA, Unpublished raw data) for 1995–2005 and Comisión Nacional de Áreas Naturales Protegidas (CONANP) & Grupo Tortuguero de Bahía de los Ángeles (Unpublished raw data) for 2005–2018. We ensured direct compatibility with LEK data by standardizing to one 12-h in-water set for a single 100m net (Seminoff et al. 2003) and analyzing mean annual CPUE values (Supporting Information: 1; Figure S1; Tables S1, S2).

### 2.4 Analyses

#### Multiple Imputation by Chained Equations (MICE)

We evaluated CPUE trends for two distinct periods: Commercial Fishing (1952–1983) and Conservation (1978–2018) (Figure 2). For the Conservation period, we appended LEK values from 1978–1982 to the monitoring dataset and interpolated values in the temporal gap between the end of commercial fishing (1983) and the start of in-water monitoring (1995). This is justified, as conservation efforts began in the late 1970s and increased over time (Figure 2) (Márquez 1996; Seminoff et al. 2008) (Supporting Information: 3).

We ran multiple imputation by chained equations (MICE) with the *mice* package *in R 4.0.4* (van Buuren & Groothuis-Oudshoorn 2011) to handle the high percentage of missing values (Commercial Fishing = 48%, Conservation = 42%). This systematic missing data method is robust for scenarios with up to 75% missingness (Takahashi 2017) (Figure 3). We assumed that values were missing at random (i.e., missingness was not systematically related to catch rates) (Thurstan et al. 2014).

**Figure 3:**
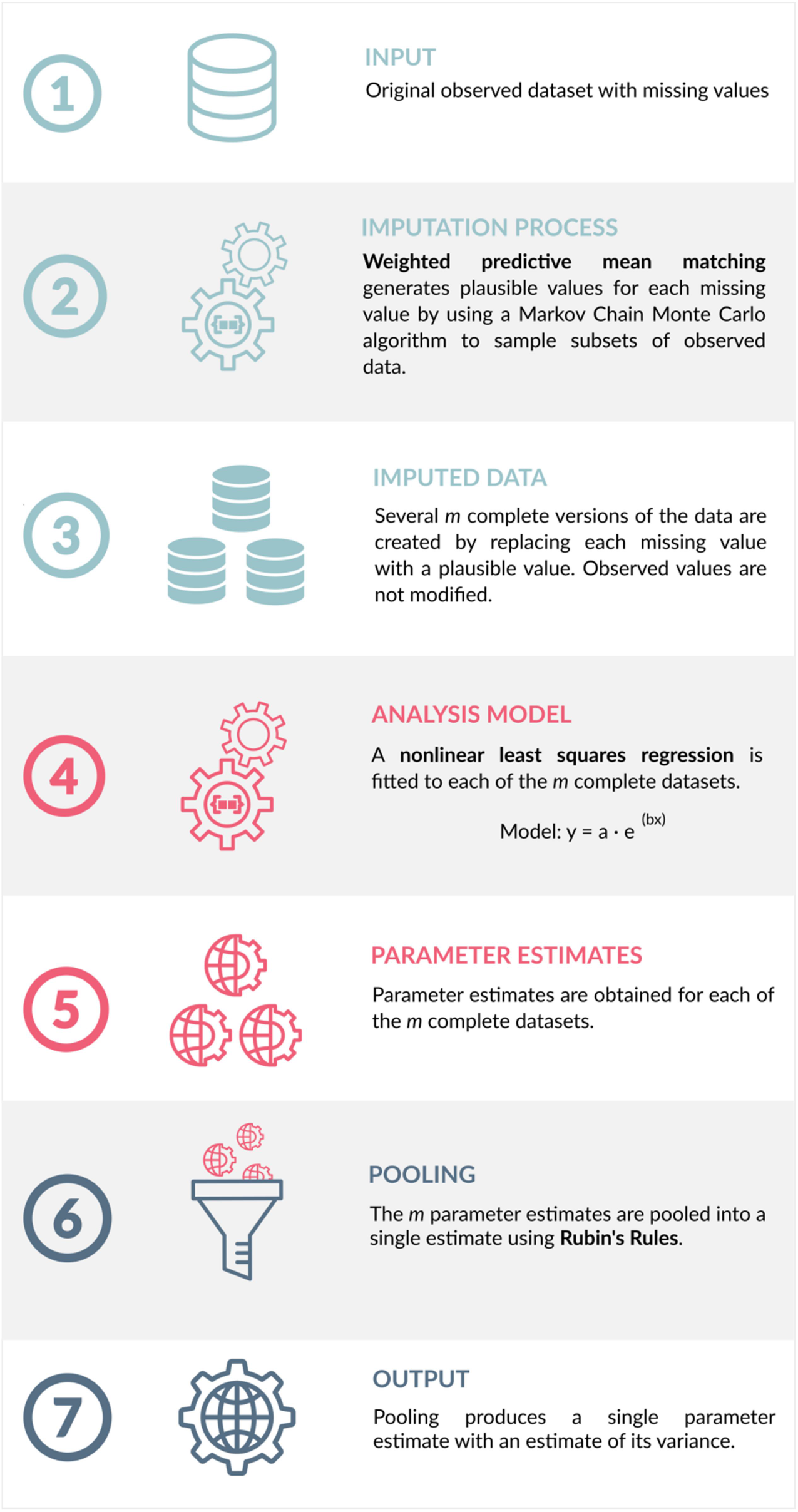
Diagram of methodological pipeline used for Multiple Imputation by Chained Equations (MICE). Models and rulesets are shown in bold.

Each missing annual value was replaced with a plausible value generated using a Markov Chain Monte-Carlo algorithm to sample subsets of observed values (van Buuren & Groothuis-Oudshoorn 2011) (Supporting Information: 4.1). We generated *m* complete datasets, equivalent to the percentage of missing values for each phase, by running 1500 iterations (Bodner 2008) (Figures S2, S3). Observed values were retained.

We fitted each of the *m* complete datasets separately to the model:

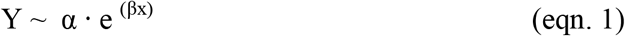

Where *Y* is the response variable (CPUE), *x* is the independent variable (year), and *α* and *β* are fitted constants. MICE does not generate a singular regression. Instead, the results of the *m* fitted models are pooled using Rubin’s Rules (Dong & Peng 2013) to give parameter estimates with standard errors that (i) describe the uncertainty of imputed missing data; (ii) account for variance within and between imputed models; and (iii) are unbiased and have valid statistical properties (van Buuren & Groothuis-Oudshoorn 2011) (Figure 3) (Supporting Information: 4.2, 4.3; Table S3).

We ratified validity through residual analysis (e_i_ ~ N(0, σ^2^)) (Nguyen et al. 2017) (Supporting Information: 4.4). We used an *ad hoc* method to (i) visualize a pooled trend line (broadly equivalent to regression line), showing pooled predicted values across all *m* multiply imputed models, and (ii) draw 95% Confidence Intervals for the pooled trend line using Rubin’s Rules (Dong & Peng 2013; Nguyen et al. 2017) (Supporting Information: 4.5).

#### Population growth and size distribution

We calculated population growth rates during Commercial Fishing (1953–1982) and Conservation (1978-2018) (Supporting Information: 5). Curved carapace length (CCL) size distributions from monitoring data were converted to life stages based on mean nester size at Colola (82.0 cm CCL) (Seminoff et al. 2015); i.e. adults > 82.0. We used Mann-Whitney U tests (α = 0.05) to compare mean annual catch rates in the LEK and monitoring datasets (Supporting Information: 6, 7), and to compare size and life stage composition over time (Period 1: 1995–2005; Period 2: 2009–2018).

## Results

Our results strongly suggest increasing green turtle abundance after ~40 years of conservation measures and ~30 years of full legal protection (Figure 4a). Both processes, Commercial Fishing and Conservation, are described by a nonlinear model (eqn. 1). The Commercial Fishing period has a high α and negative β (α = 24.271, β = −0.820; R^2^ = 0.845) while during the Conservation period a low α and positive β (α = 0.002, β = 0.136; R^2^ = 0.711), both with robust residuals and good fits (Figure 5; Table 1).

**Table 1:** Results of Multiple Imputation by Chained Equations (MICE) analysis with non-linear model. Bold type indicates significant results at α = 0.05. Parameter estimates, confidence intervals, standard error, and R^2^ values were pooled using Rubin’s Rules to account for uncertainty of the missing data and variance within and between the *m* imputed models (Dong & Peng, 2013). 95% confidence intervals for R^2^ values are shown in brackets. Pooled degrees of freedom are included to account for the effects of missing data (Supporting Information:4.6) (van Buuren & Groothuis-Oudshoorn, 2011). See also Figure 5.

| Parameter | Pooled estimate | Pooled 95% C.I. | Pooled std. error | P-value | Pooled *df* | Pooled *R*<sup>2</sup> |
| --- | --- | --- | --- | --- | --- | --- |
| *Commercial Fishing (1952-1982; LEK data)* |  |  |  |  |  |  |
| *Model: $y \sim \alpha^{(\beta \cdot x)}$; *df* (complete data) = 29; *m* = 48; $e \sim N(0, \sigma^2)$ for mean residuals* |  |  |  |  |  |  |
| $\alpha$ | 24.271 | [19.669 - 28.873] | 2.189 | < **0.01** | 17.800 | |
| $\beta$ | -0.0820 | [-0.101 to -0.0628] | 0.00928 | < **0.01** | 24.060 | 0.845 [0.692-0.926] |
| *Conservation (1978-2018; LEK and Monitoring data\*)* |  |  |  |  |  |  |
| *Model: $y \sim \alpha^{(\beta \cdot x)}$; *df* (complete data) = 38; *m* = 42; $e \sim N(0, \sigma^2)$ for mean residuals* |  |  |  |  |  |  |
| $\alpha$ | 0.0023 | [-0.0122 - 0.0168] | 0.00703 | 0.746 | 25.325 | |
| $\beta$ | 0.136 | [ 0.0760 - 0.196] | 0.0286 | < **0.01** | 17.376 | 0.711 [0.364-0.890] |

\* LEK values from 1978-1982 were appended to the monitoring dataset to allow for interpolation of values in the temporal gap between LEK and monitoring datasets.

**Figure 4:**
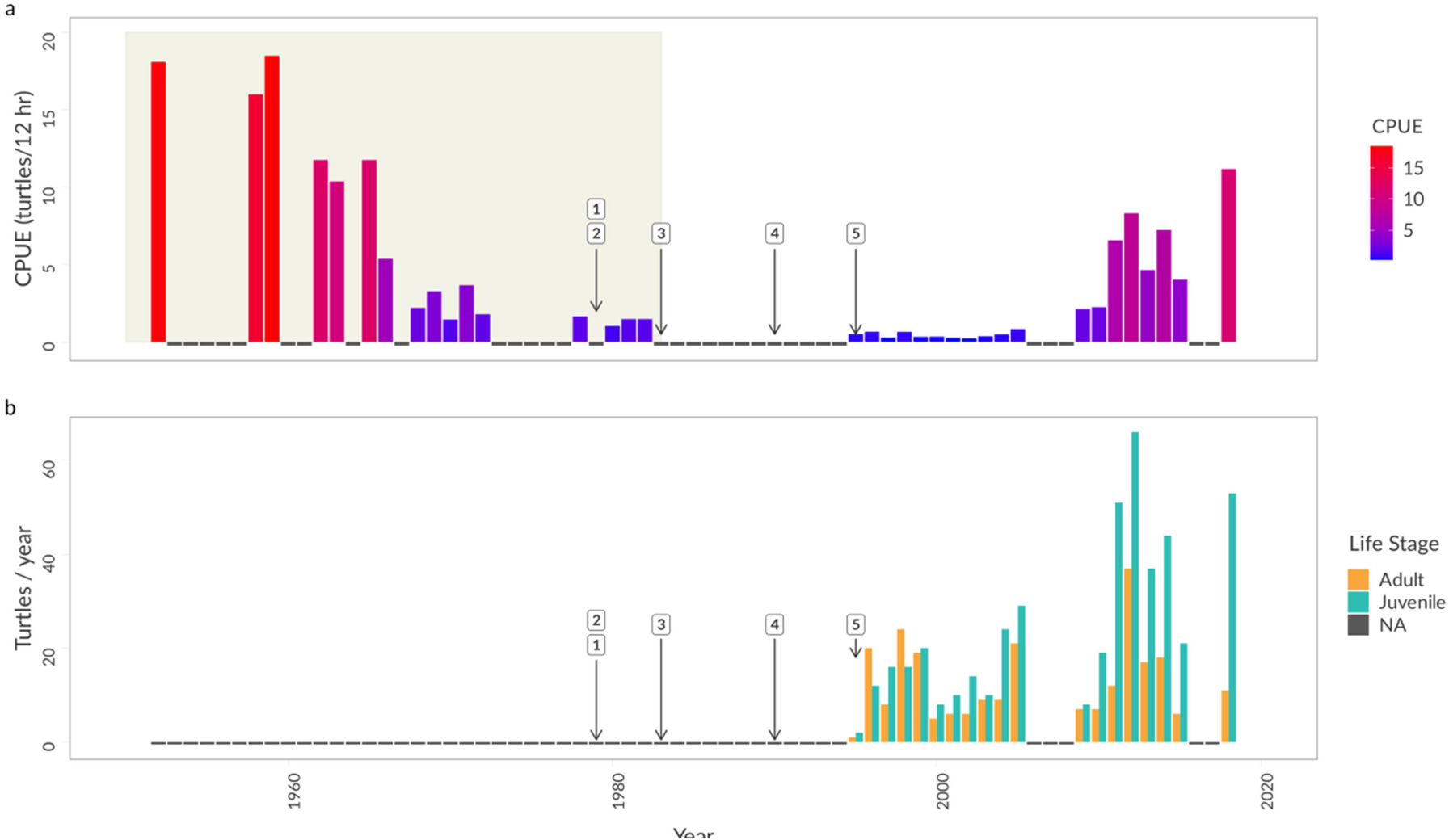
Long-term trends in mean annual catch-per-unit-effort (CPUE) (a) and turtle life stage distribution (b). Note difference in the y-axis scale. Annotations show key events in sea turtle conservation and management: (1) start of permanent sea turtle research efforts at Bahía de los Ángeles; (2) start of nesting beach protection at Colola, Michoacán; (3) suspension of green turtle fishing permits by the Federal Government; (4) permanent ban on all sea turtle captures in Mexico; (5) start of in-water monitoring at Bahía de los Ángeles. Panel (a) shows mean annual CPUE values from 1952-2018. Shaded area shows data derived from Local Ecological Knowledge. Panel (b) shows total captures per year in scientific monitoring for adults (Curved Carapace Length, CCL > 82.0 cm) and juveniles (CCL ≤ 82.0 cm). Size at maturity is based on mean size of nesting females at Colola (Figueroa et al. cited in Seminoff et al. 2015).

**Figure 5:**
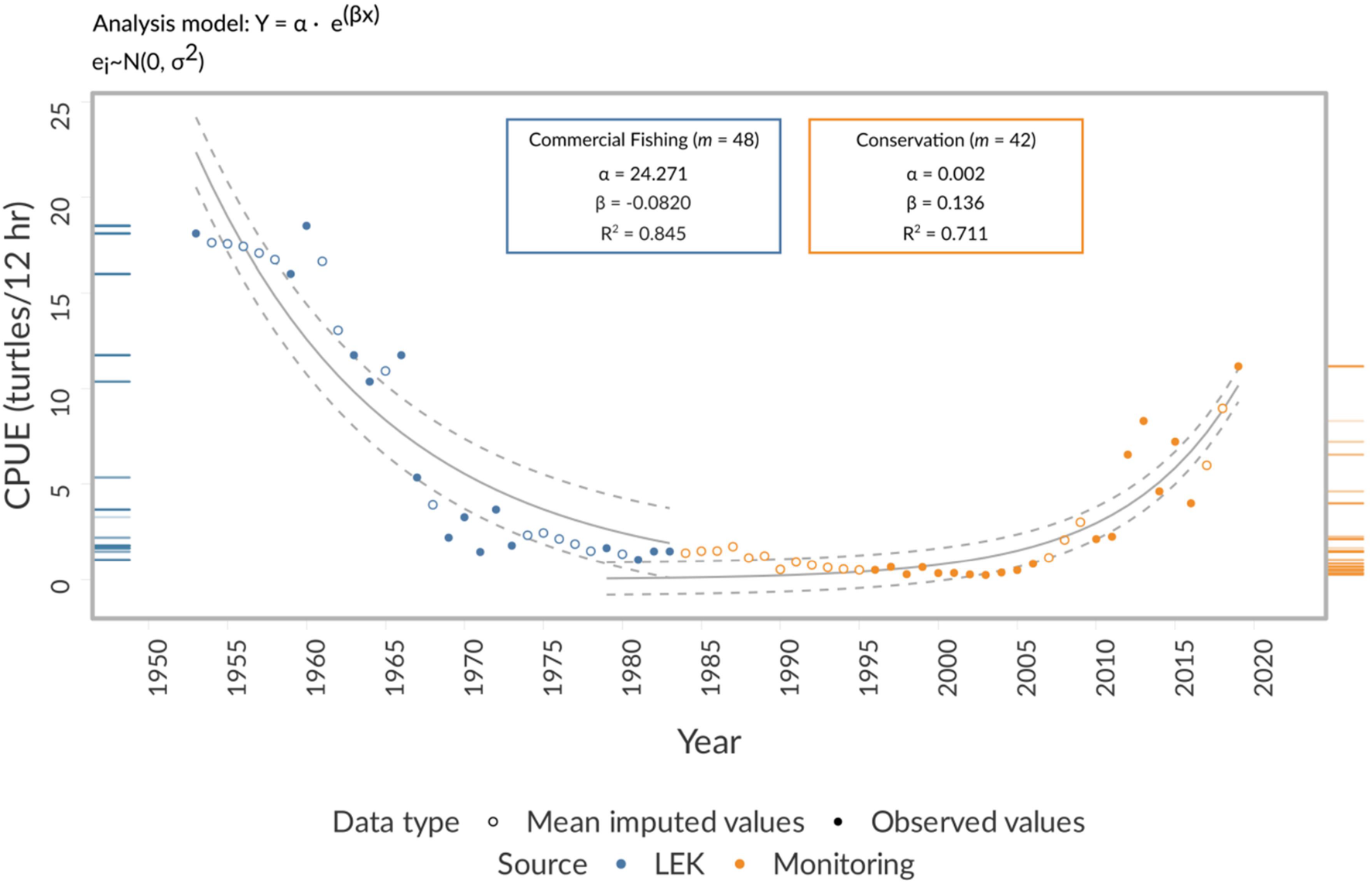
Trends in catch-per-unit-effort (CPUE) for 1952–2018 using Multiple Imputation by Chained Equations (MICE). Data points show mean value for each year. Pooled parameter estimates and R^2^ values are shown for the Commercial Fishing phase (1952– 1983; blue box) and the Conservation phase (1978–2018; orange box). Solid trend line shows pooled predicted values across all *m* imputed models, and dashed lines show 95% Confidence Intervals for the pooled trend line. Standard errors were pooled according to Rubin’s Rules to account for within-model and between-model variance (Dong & Peng 2013) (Supporting Information: 4.5). Marginal rug plots show density distributions of imputed values for Commercial Fishing (blue rug plot, n = 630) and Conservation (orange rug plot, n = 816) (see also Figures S2 and S3). Pooled 95% Confidence Intervals for parameter estimates and R^2^ values are reported in Table 1.

The BLA population declined at a rate of 8.4% annually during commercial fishing, which contrasts with the 4.8% annual increase during the conservation phase (Table S4), emphasizing that declines occurred 75% faster than increases. Notably, there is a prolonged latency period between the initial implementation of conservation measures (1979), initial signs of increase (~2000), and the significant recovery phase after 2011. Despite the clear upward trend, maximum CPUE during scientific endeavor (11.2 turtles/12hr; 2018) represents ~60% of the maximum CPUE in the commercial fishery (18.5 turtles/12hr; 1959) (Figures 4a, 5). Furthermore, median CPUE is significantly lower in scientific monitoring (Median = 0.66) than commercial fishing (Median = 3.47) (Mann-Whitney U = 232, p <0.05, 95% C.I. [0.81–7.75]) (Tables S2, S5). Thus, although abundance levels in BLA are near those of the mid-1960s, they still remain below the historical baseline.

### Life stage and size distribution (1995–2018)

Abundance increases after 2011 coincide with shifts toward a higher proportion of juvenile turtles (Figure 4b). Period 2 (2009–2018; 72.22% juveniles) shows a strong juvenile bias compared with Period 1 (1995–2005) (55.71% juveniles). Furthermore, median CCL was significantly smaller in Period 2 (Median = 75.5) than Period 1 (Median = 80.8) (Mann-Whitney U = 71406, p <0.001, 95% C.I. [1.99–5.20]) (Tables S6, S7). These patterns suggest juvenile recruitment drives population growth and are consistent with sea turtles’ prolonged somatic growth rates, which require decades of hatchling production before juvenile recruitment into foraging habitats is observed (Seminoff & Shanker 2008).

## Discussion

The robust positive trend for foraging populations at BLA shows that conservation measures work, but require prolonged time frames. While intensive, technologically efficient commercial fishing —even by a small artisanal fleet— can have severe impacts in a relatively short time (Peckham et al. 2007), population recovery requires decades of sustained protection across habitats. In our case, whereas collapse occurred in ~20 years, significant population growth was only possible after ~40 years of conservation. This pattern of fast decline and slow population growth is consistent with other long-lived marine taxa, including large sharks, sirenids, and cetaceans (cf. Chaloupka et al. 2008; McClenachan et al. 2012).

Our results further demonstrate that for depleted long-lived, highly migratory species like sea turtles, spatially and temporally widespread conservation measures — nesting beach and habitat protection, by-catch regulation, and protection from unsustainable use of meat or eggs— are pre-requisites for positive results (Chaloupka et al. 2008). The BLA foraging population shows encouraging trends. Nonetheless, current local abundance remains significantly below baseline levels. This result is consistent with the consensus among senior fishers that green turtles are abundant but below numbers observed when they were young harpooners (Early-Capistrán et al. 2020a). Thus, continued conservation measures are required for the population to return to historical levels.

### The need for integrative analysis

The evaluation of green turtle conservation status is complicated by the lack of knowledge about fundamental parameters, including maturation age, life-stage duration, and migratory and demographic connectivity (Casale & Heppell 2016; Seminoff & Shanker 2008). Holistic assessment is beyond the scope of this paper. However, population trajectories at BLA largely coincide with trends at the Colola index nesting site. The Colola coastline was largely uninhabited until the 1950s. Settlements grew as market demand for sea turtle products surged in the early 1970s, and ~70,000 eggs/night were harvested commercially until nesting beach protection began in 1979 (Delgado-Trejo & Alvarado Díaz 2012). Like BLA’s foraging population, the rookery at Colola has grown substantially since ~2010: nesting rates increased 508% from 1982 (3,383 nests/year) to 2015 (15,196 nests/year). Nevertheless, the degree of relative increase remains unclear due to the lack of pre-exploitation baseline data for Colola (Delgado-Trejo 2016).

Synchronous over-exploitation across life-stages and habitats from 1960-1980 likely contributed to steep population declines by simultaneously decreasing adult survivorship, hatchling production, and juvenile recruitment (cf. Seminoff & Shanker 2008). Reversing the negative trends was achieved by concurrent conservation measures across the species’ range in Mexico (Figure 2). Decades of fishing bans and nesting beach protection have generated a positive feedback loop of increased survivorship and recruitment across life stages and habitats. Comparisons of trends between nesting and foraging habitats will provide insight to be evaluated in future research (Van Houtan et al. 2014).

### Implications for conservation

Our results suggest a robust causal relationship between abundance trends and key events in conservation and management policies (Figure 4a). BLA’s foraging population shows an encouraging trend thanks to decades of conservation measures enacted over broad spatial scales. Future research agendas will benefit from integrating the effect of climatic fluctuations, which are drivers of sea turtle population dynamics including hatchling production, sex determination, juvenile recruitment, foraging success, and the timing and frequency of reproduction (Patrício et al. 2021). Threats from climate change are more difficult to mitigate than direct human activity, making tangible conservation efforts increasingly challenging (Mazaris et al. 2017).

While efforts at national, international, and boundary-wide scales are necessary for effective green turtle conservation, data from foraging areas are crucial for informing local management (Bjorndal et al. 2005; Broderick et al. 2006). Likewise, multiple forms of expertise —including locally-grounded collaboration— are essential for creating truly diverse and inclusive approaches to conservation (cf. Carman & González Carman 2020). Our results were only possible thanks to long-term, collaborative efforts with the BLA community, as LEK was indispensable for establishing baseline levels, determining local recovery targets, and evaluating current population status. Importantly, LEK must be recognized for its inherent value, and become integral to conservation policy and practice. As such, any future conservation initiatives —i.e., establishing local habitat protection areas, by-catch reduction in commercial fisheries, or locally grounded strategies to prevent unregulated fishing that impact green turtles and their habitats— must be determined by local communities, with self-determination as the guiding principle of scientific collaboration (cf. Mawyer & Jacka 2018).

### Global perspectives and future challenges

Patterns of historical green turtle abundance, decline from overfishing, and growth thanks to conservation efforts are documented throughout the Pacific Rim, Atlantic, Caribbean, and Indo-Pacific regions (Broderick et al. 2006; Mazaris et al. 2017). However, climate change will pose new and increasing challenges (cf. Patrício et al. 2021). LEK-based approaches will become increasingly relevant as sea turtle populations grow and cease to be mere conservation targets (cf. Christianen et al. 2021), particularly considering that conservation conflicts have arisen when management frameworks fail to account for the sea turtles’ cultural importance (cf. Barrios-Garrido et al. 2018). To be successful, future conservation and policy measures for migratory marine species must integrate international and basin-wide approaches built upon locally-grounded efforts, integrating the diverse peoples and worldviews linked to the oceans (Vierros et al. 2020). LEK, accumulated by people living with and from the sea, is indispensable for comprehending long-term change and building sustainable futures.

## Acknowledgements and data

We thank the community of Bahía de los Ángeles and Posgrado en Ciencias del Mar y Limnología, UNAM. Research was funded by CONACYT doctoral fellowship (#289695) to M.M.E.C.. We thank CONANP and Grupo Tortuguero de Bahía de los Ángeles for monitoring data. We thank R. Ávalos, J. Candela, and V. Koch. The data that support the findings of this study include data from CONANP, but restrictions apply to the availability of these data, which were used under license for the current study, and so are not publicly available. Data are however available from the authors upon reasonable request and with permission of CONANP. Code was implemented in R and is available upon request.

## Ethical approval

Research was approved by the Bioethics Committee of the Centro de Investigación Científica y de Educación Superior de Ensenada (Approval Number 2S.3.1).

## References

Barrios-Garrido, H.A., Palmar, J., Wildermann, N., Rojas Izales, D., Diedrich, A. & Hamann, M. (2018). Marine turtle presence in the traditional pharmacopoeia, cosmovision, and beliefs of Wayuú indigenous people. Chelonian Conservation and Biology, 17, 177.

Bjorndal, K.A., Bolten, A.B. & Chaloupka, M.Y. (2005). Evaluating trends in abundance of immature green turtles, *Chelonia mydas*, in the greater Caribbean. Ecological Applications, 15, 304–314.

Bodner, T.E. (2008). What improves with increased nissing data imputations? Structural Equation Modeling: A Multidisciplinary Journal, 15, 651–675.

Broderick, A.C., Frauenstein, R., Glen, F., Hays, G.C., Jackson, A.L., Pelembe, T., Ruxton, G.D. & Godley, B.J. (2006). Are green turtles globally endangered? Global Ecology and Biogeography, 15, 21–26.

van Buuren, S. & Groothuis-Oudshoorn, K. (2011). MICE : multivariate imputation by chained equations in R. Journal of Statistical Software, 45.

Carman, M. & González Carman, V. (2020). Going beyond diverse worldviews for conservation: response to Kohler et al. Conservation Biology, 34, 286–288.

Casale, P. & Heppell, S. (2016). How much sea turtle bycatch is too much? A stationary age distribution model for simulating population abundance and potential biological removal in the Mediterranean. Endangered Species Research, 29, 239–254.

Chaloupka, M., Bjorndal, K.A., Balazs, G.H., Bolten, A.B., Ehrhart, L.M., Limpus, C.J., Suganuma, H.S., Troëng, S. & Yamaguchi, M. (2008). Encouraging outlook for recovery of a once severely exploited marine megaherbivore. Global Ecology and Biogeography, 17, 297–304.

Christianen, M.J.A., van Katwijk, M.M., van Tussenbroek, B.I., Pagès, J., Ballorain, K., Kelkar, N., Arthur, R. & Alcoverro, T. (2021). A dynamic view of seagrass meadows in the wake of successful green turtle conservation. Nature Ecology and Evolution.

Delgado-Trejo, C. (2016). Recovery of the black sea turtle in Michoacan, Mexico: final report to the U.S. Fish and Wildlife Service (Technical report). U.S. Fish and Wildlife Service and Universidad Michoacana San Nicolás Hidalgo, Morelia, Mexico.

Delgado-Trejo, C. & Alvarado Díaz, J. (2012). Current conservation status of the black sea turtle in Michoacán, Mexico. In: Sea Turtles of the Eastern Pacific: Advances in Research and Conservation (eds. Seminoff, J.A. & Wallace, B.P.). University of Arizona Press, Tucson, pp. 263–278.

Diario Oficial de la Federación. (1990). Acuerdo por el que se establece veda para las especies y subespecies de tortuga marina en aguas de jurisdicción Federal del Golfo de México y Mar Caribe, así como en las del Océano Pacífico, incluyendo el Golfo de California. *DOF 31/05/1990*.

Dong, Y. & Peng, C.-Y.J. (2013). Principled missing data methods for researchers. SpringerPlus, 2, 222.

Dutton, P.H., LeRoux, R.A., LaCasella, E.L., Seminoff, J.A., Eguchi, T. & Dutton, D.L. (2019). Genetic analysis and satellite tracking reveal origin of the green turtles in San Diego Bay. Mar Biol, 166, 3.

Early-Capistrán, M.-M., Sáenz-Arroyo, A., Cardoso-Mohedano, J.-G., Garibay-Melo, G., Peckham, S.H. & Koch, V. (2018). Reconstructing 290 years of a data-poor fishery through ethnographic and archival research: The East Pacific green turtle (*Chelonia mydas*) in Baja California, Mexico. Fish Fish, 19, 57–77.

Early-Capistrán, M.-M., Solana-Arellano, E., Abreu-Grobois, F.A., Narchi, N.E., Garibay-Melo, G., Seminoff, J.A., Koch, V. & Saenz-Arroyo, A. (2020a). Quantifying local ecological knowledge to model historical abundance of long-lived, heavily-exploited fauna. PeerJ, 8, e9494.

Early-Capistrán, M.-M., Solana-Arellano, E., Abreu-Grobois, F.A., Narchi, N.E., Garibay-Melo, G., Seminoff, J.A., Koch, V. & Saenz-Arroyo, A. (2020b). Quantitative datasets and R Code: Quantifying local ecological knowledge to model historical abundance of long-lived, heavily-exploited fauna. PeerJ, 8, e9494.

Game, E.T., Schwartz, M.W. & Knight, A.T. (2015). Policy relevant conservation science. Conservation Letters, 8, 309–311.

Garibaldi, A. & Turner, N. (2004). Cultural Keystone Species: Implications for ecological conservation and restoration. E&S, 9, art1.

IUCN. (2019). The IUCN Red List of Threatened Species [WWW Document]. IUCN Red List of Threatened Species. *Version 2019-1.* URL https://www.iucnredlist.org/en

Koch, V. (2013). 12 años de monitoreo de la tortuga negra (Chelonia mydas) en zonas de alimentación y crianza en el Noroeste de México (Technical report). Grupo Tortuguero de las Californias A.C., La Paz, B.C.S., Mexico.

Lee, L.C., Thorley, J., Watson, J., Reid, M. & Salomon, A.K. (2018). Diverse knowledge systems reveal social-ecological dynamics that inform species conservation status. Conservation Letters, e12613.

Márquez, R. (1996). Las tortugas marinas y nuestro tiempo. La ciencia para todos. Fondo de Cultura Económica, Mexico City.

Mawyer, A. & Jacka, J.K. (2018). Sovereignty, conservation and island ecological futures. Envir. Conserv., 45, 238–251.

Mazaris, A.D., Schofield, G., Gkazinou, C., Almpanidou, V. & Hays, G.C. (2017). Global sea turtle conservation successes. Science Advances, 3, e1600730.

McClenachan, L.E., Ferretti, F. & Baum, J.K. (2012). From archives to conservation: why historical data are needed to set baselines for marine animals and ecosystems: From archives to conservation. Conservation Letters, 5, 349–359.

Nguyen, C.D., Carlin, J.B. & Lee, K.J. (2017). Model checking in multiple imputation: an overview and case study. Emerging Themes in Epidemiology, 14.

Patrício, A., Hawkes, L., Monsinjon, J., Godley, B. & Fuentes, M. (2021). Climate change and marine turtles: recent advances and future directions. Endang. Species. Res., 44, 363–395.

Peckham, S.H., Diaz, D.M., Walli, A., Ruiz, G., Crowder, L.B. & Nichols, W.J. (2007). Small-scale fisheries bycatch jeopardizes endangered Pacific loggerhead turtles. PLoS ONE, 2, e1041.

Poe, M.R., Norman, K.C. & Levin, P.S. (2014). Cultural dimensions of socioecological systems: Key connections and guiding principles for conservation in coastal environments. Conservation Letters, 7, 166–175.

Sáenz-Arroyo, A. & Revollo-Fernández, D. (2016). Local ecological knowledge concurs with fishing statistics: An example from the abalone fishery in Baja California, Mexico. Marine Policy, 71, 217–221.

Selgrath, J.C., Gergel, S.E. & Vincent, A.C.J. (2018). Shifting gears: Diversification, intensification, and effort increases in small-scale fisheries (1950-2010). PLOS ONE, 13, e0190232.

Seminoff, J.A., Allen, C.D., Balazs, G., Dutton, P.H., Eguchi, T., Haas, H.L., Hargrove, S., Jensen, M., Klemm, D.L., Lauritsen, A.M., MacPherson, S.L., Opay, P., Possardt, E.E., Pultz, S., Seney, E., Van Houtan, K.S. & Waples, R.S. (2015). Status review of the green turtle (Chelonia mydas) under the Endangered Species Act (NOAA Technical Memorandum No. NOAA-TM-NMFS-SWFSC-539). NOAA, San Diego, USA.

Seminoff, J.A., Jones, T.T., Resendiz, A., Nichols, W.J. & Chaloupka, M.Y. (2003). Monitoring green turtles (*Chelonia mydas*) at a coastal foraging area in Baja California, Mexico: multiple indices to describe population status. Journal of the Marine Biological Association of the UK, 83, 1355–1362.

Seminoff, J.A., Reséndiz-Hidalgo, A., Jiménez de Reséndiz, B., Nichols, W.J. & Todd-Jones, T. (2008). Tortugas marinas. In: Bahía de los Ángeles: recursos naturales y comunidad: línea base 2007 (eds. Danemann, G. & Ezcurra, E.). Secretaría de Medio Ambiente y Recursos Naturales ; San Diego Natural History Museum, Tlalpan, México D.F.; San Diego, Calif., pp. 457–494.

Seminoff, J.A. & Shanker, K. (2008). Marine turtles and IUCN Red Listing: A review of the process, the pitfalls, and novel assessment approaches. Journal of Experimental Marine Biology and Ecology, 356, 52–68.

Takahashi, M. (2017). Statistical inference in missing data by MCMC and non-MCMC multiple imputation algorithms: Assessing the effects of between-imputation iterations. Data Science Journal, 16, 37.

Thurstan, R.H., Campbell, A.B. & Pandolfi, J.M. (2014). Nineteenth century narratives reveal historic catch rates for Australian snapper (*Pagrus auratus*). Fish and Fisheries, 2–16.

Van Houtan, K.S., Hargrove, S.K. & Balazs, G.H. (2014). Modeling Sea Turtle Maturity Age from Partial Life History Records. Pacific Science, 68, 465–477.

Vierros, M.K., Harrison, A.-L., Sloat, M.R., Crespo, G.O., Moore, J.W., Dunn, D.C., Ota, Y., Cisneros-Montemayor, A.M., Shillinger, G.L., Watson, T.K. & Govan, H. (2020). Considering Indigenous Peoples and local communities in governance of the global ocean commons. Marine Policy, 119, 104039.

