## Supporting Information for "Integrating ecological monitoring and local ecological knowledge to evaluate conservation outcomes"

**Integrating ecological monitoring and local ecological knowledge to evaluate conservation outcomes (bioRxiv pre-print)**

Michelle María Early-Capistrán[1], Elena Solana-Arellano[2], F. Alberto Abreu-Grobois[3], Gerardo Garibay-Melo [4], Jeffrey A. Seminoff [5], Andrea Sáenz-Arroyo [6], Nemer E. Narchi[7]

[1] Posgrado en Ciencias del Mar y Limnología, Universidad Nacional Autónoma de México, Mexico City, Mexico
[2] Departamento de Ecología Marina, Centro de Investigación Científica y Educación Superior de Ensenada, Ensenada, Baja California, Mexico.
[3] Unidad Académica Mazatlán, Instituto de Ciencias del Mar y Limnología, Universidad Nacional Autónoma de México, Mazatlán, Sinaloa, Mexico.
[4] Posgrado en Manejo de Ecosistemas de Zonas Áridas. Universidad Autónoma de Baja California, Ensenada, B.C., Mexico.
[5] NOAA – Southwest Fisheries Science Center, La Jolla, California, U.S.
[6] Departamento de Conservación de la Biodiversidad. El Colegio de la Frontera Sur (ECOSUR), San Cristóbal de Las Casas, Chiapas, Mexico
[7] CoLaboratorio de Oceanografía Social/Centro de Estudios de Geografía Humana, El Colegio de Michoacán - Sede La Piedad, La Piedad, Michoacán, Mexico

Early-Capistrán et al. (2021) Pre-print

Early-Capistrán et al. (2021) Pre-print

#### 1. Monitoring procedures

During scientific monitoring, turtles were captured using entanglement nets with the same depth (8m) and mesh size (0.8m) as those used by commercial turtle fishers. Nets were set and checked every 0.5-12hr, and captured turtles were measured, weighed, and released (Seminoff et al. 2003; Santacruz López 2012). Nets are set for periods of 12-24 hr, and monitoring occurs monthly, with some variation due to weather conditions, logistics, and funding availability (Seminoff et al. 2003; Santacruz López 2012).

#### 2. Mean values for analyses

Analyses of commercial green turtle fishing revealed that high CPUE events occurred throughout the chronology despite overall declines in abundance (hyper-stability), as expert fishers consistently targeted high-density locations or aggregations of turtle by relying on detailed empirical knowledge of green turtle behavior and local environmental and oceanographic conditions (Early-Capistrán et al. 2020). Thus, methods for compiling LEK data focused on estimating central trends rather than high catch events, and we used mean, standardized CPUE values to obtain representative estimates (Early-Capistrán et al. 2020). We used the same approach to evaluate monitoring data, (i) to maintain consistency throughout the time series, and (ii) to reduce the effects of high variability in catch rates, likely due to green turtle grouping behavior (Figure S1).

Early-Capistrán et al. (2021) Pre-print

#### 3. Commercial fishing and Conservation phases

For MICE analyses and calculations of population growth rates, we grouped data into two phases: Commercial Fishing (1952-1983) and Conservation (1978-2018) (Figure 2). For the Conservation phase, we appended LEK values from 1978-1982 to the monitoring dataset. These values correspond to the fishery collapse stage (Figure 2) (Early-Capistrán et al. 2020). Long-term conservation and research efforts also began in the late 1970s in response to diminishing green turtle populations (Seminoff et al. 2008). We consider this to be an adequate modification, as conservation and research efforts began in the late 1970s and regulation increased over time, culminating in the permanent ban on all sea turtle capture and use in Mexico, declared in 1990 (Figure 2). Integrating LEK and monitoring values for the Conservation phase allowed us to interpolate values in the temporal gap between the end of commercial fishing (1983) and the start of in-water monitoring (1995). The Commercial Fishing phase is described with the LEK dataset, to which no data were appended.

Early-Capistrán et al. (2021) Pre-print

#### 4. Analyses with Multiple Imputation by Chained Equations (MICE)

##### 4.1 Imputation model selection

We selected the imputation model by comparing three methods: predictive mean matching (pmm), weighted predictive mean matching (wpmm), and Bayesian linear models (van Buuren 2018). Both pmm and wpmm implement a Markov Chain Monte-Carlo algorithm to sample subsets of observed values (van Buuren 2018). For each imputation model, we generated *m* datasets equivalent to the percentage of missing data (Bodner 2008). We first observed imputed values directly, and discarded Bayesian linear models as they generated implausible negative values. We then conducted descriptive evaluation of (i) plots of the density distribution of observed and imputed values; (ii) one-dimensional scatterplots of observed and imputed values; and (iii) two-dimensional scatter plots (cpue ~ year) of observed and imputed values (van Buuren & Groothuis-Oudshoorn 2011; van Buuren 2018).

We fitted datasets imputed with pmm and wpmm to the analysis model (eqn. 1) and ran multiply imputed repeated analyses separately. We compared pooled parameter, $R^2$ values, and residuals ($e_i \sim N(0, \sigma^2)$) between imputation models, and also compared these results with the model fitted analysis for the observed values only. The results suggest that values from weighted predictive mean matching (wpmm) were more plausible than those obtained from predictive mean matching (pmm), as they had greater concordance with observed values, pooled parameter estimates were closer to estimates from observed values, and pooled residuals were robust. We selected wpmm as the imputation model for both LEK and monitoring datasets. Additionally, wpmm is robust to non-normal data distributions (Jia 2016).

**4.2 Analysis model selection**

We used a nonlinear model to describe both Commercial Fishing and Conservation processes:

$$Y \sim \alpha \cdot e^{(\beta x)} \qquad \text{(eqn. S1)}$$

Where $Y$ is the response variable (CPUE), $\alpha$ and $\beta$ are fitted constants, and $x$ is the independent variable (year), which was serialized for all analyses. We then used *R 4.0.4* to run

Early-Capistrán et al. (2021) Pre-print

nonlinear least squares regression (NLR) (Table S3) (Early-Capistrán et al. 2020). Model performance was ratified with residual analyses ($e_i \sim N(0, \sigma^2)$).

##### 4.3 Starting value selection

We used a quasi-Newton method to optimize starting values for NLR, both for observed values and for the analysis model in MICE. We obtained initial approximate starting values by linearizing the nonlinear analysis model (eqn. S1) to the form:

$$\ln(Y) \sim \ln(\alpha) + \beta x \qquad \text{(eqn. S2)}$$

We ran a linear regression with observed values, extracted parameter values, and transformed $\ln(\alpha)$ to $\alpha$. These values were used as starting values for NLR, which were then run iteratively by extracting parameter values and plugging them back in as starting values for the next iteration until parameter estimates stabilized. With MICE, we used parameter estimates from the *m* pooled models as new starting values.

##### 4.4 Residual analysis of MICE models

We evaluated model performance through residual analyses ($e_i \sim N(0, \sigma^2)$) with two complementary approaches. We first extracted and analyzed residuals from each of the *m* model fittings separately (Nguyen et al. 2017). We then averaged residuals across all *m* models to ratify the fit of the pooled model.

Early-Capistrán et al. (2021) Pre-print

##### 4.5 Confidence intervals for MICE models

Graphic representation of results generated by MICE represents a unique challenge, as this method generates *m* "complete" datasets in which missing data are replaced by plausible simulated values and, afterward, the analysis model (eqn. 1 in main text) is fitted separately to each of the *m* datasets and then pooled using Rubin's Rules to generate pooled parameter estimates with standard errors that (i) account for variance within and between imputed models, and (ii) allow for the uncertainty of the missing data (Dong & Peng 2013; Nguyen et al. 2017). If performed under appropriate conditions, pooling preserves the relations between the data, preserve the uncertainty about these relations, and generate results that are unbiased and have valid statistical properties (van Buuren & Groothuis-Oudshoorn 2011; van Buuren 2018). However, MICE does not generate a singular model or regression, but rather describes the distribution of the parameters generated by the *m* multiply imputed models (Bodner 2008; Nguyen et al. 2017).

We used an *ad hoc* method to (i) visualize a trend line (broadly equivalent to a regression line) that reflects the pooled predicted values across all *m* multiply imputed models, (ii) pool values for the upper and lower bounds according to Rubin's Rules to account for both within-model and between-model variance across the *m* multiply imputed models, and (iii) draw 95% confidence intervals describing the upper and lower bounds for all points of the pooled trend line (Dong & Peng 2013; Nguyen et al. 2017). The process in *R* (4.0.4) consisted of the following steps (see also Data and Code):

Early-Capistrán et al. (2021) Pre-print

- Calculate predicted values across all $m$ multiply imputed models to generate $m$ vectors of predicted values for each year
- Obtain pooled predictions by calculating the mean predicted value for each year across the $m$ multiply imputed models, to obtain the trend line.
- Calculate within-model variance for each of the $m$ multiply imputed models with the following equation:

$$Vw = \frac{1}{m}\left(\sum_{i=1}^{m} SE^2\right) \qquad \text{(eqn. S3)}$$

  - Where $m$ is the number of imputed datasets and SE is the estimated standard error of each of the imputed datasets.

- Calculate between-model variance of the $m$ multiply imputed models with the following equation:

$$Vb = \frac{\sum_{i=1}^{m}\left(\theta - \overline{\theta}\right)^2}{(m-1)} \qquad \text{(eqn. S4)}$$

  - Where $\theta$ is the parameter of interest (the mean of each of the $m$ vectors of predicted values) and $\overline{\theta}$ is the mean parameter value (in this case, the mean of the means).

- Calculate total variance with the following equation:

Early-Capistrán et al. (2021) Pre-print

$$Vtotal = Vw + Vb + 1/Vb \qquad \text{(eqn. S5)}$$

- Calculate the pooled standard error with the following equation:

$$SEpooled = \sqrt{V}total \qquad \text{(eqn. S6)}$$

- Calculate 95% confidence intervals of ± 1.96 standard errors from the mean, given the normal distribution of errors verified in residual analysis.

Early-Capistrán et al. (2021) Pre-print

##### 4.6 Degrees of freedom in multiple imputation

Degrees of freedom in multiple imputation must account for the effects of missing data, generating adjusted degrees of freedom for each parameter (van Buuren 2018). The *mice* package in *R* (van Buuren & Groothuis-Oudshoorn 2011) uses Barnard-Rubin's adjustment (Barnard & Rubin 1999) to calculate degrees of freedom for the pooled parameter estimates, *v*, using the degrees of freedom of a hypothetically complete dataset, $\bar{Q}$, and $\lambda$, which describes the proportion of variation due to missing data (van Buuren 2018). Here we provide a brief conceptual overview. Full mathematical and computational details are available in van Buuren (2018), Chapter 2.

Let *m* be the number of imputed datasets. Rubin's (1987) original equation for degrees of freedom is as follows:

$$v_r = \frac{(m-1)}{\lambda^2} \qquad \text{(eqn. S7)}$$

Estimated degrees of freedom for observed data which account for the effects of missing data, $v_{obs}$, are calculated as follows:

$$v_{obs} = \frac{(v_{com}+1)}{(v_{com}+3)} v_{com}(1-\lambda) \qquad \text{(eqn. S8)}$$

$v_{com}$ are the degrees of freedom of a hypothetical complete dataset, $\bar{Q}$. For a model with $k$ parameters and a sample size $n$, $v_{com}$ can be defined as $v_{com} = n - k$ (van Buuren 2018). Barnard and Rubin's (1999) adjusted degrees of freedom for use in tests with multiple imputation are calculated as follows:

$$v = \frac{v_r \cdot v_{obs}}{v_r + v_{obs}} \qquad \text{(eqn. S9)}$$

The adjusted value $v$ is always less than or equal to $v_{com}$ (van Buuren 2018). To broadly describe the effect of the variance ratios, consider that λ occupies values between 0 and 1. If λ = 0, then $v = v_{com}$; that is to say, there is no variation due to missingness. Conversely, λ = > 0.5 suggests substantial effects due to imputation (van Buuren 2018).

#### 5. Population growth rates

We calculated population growth rates for the processes of Commercial Fishing (1952-1983) and Conservation (1978-2018) by modifying the analysis model (eqn. 1) to the form:

$$N = N_0 \cdot e^{(rt)} \qquad \text{(eqn. S10)}$$

Early-Capistrán et al. (2021) Pre-print

Where $N$ is the end value, $N_0$ is the starting value, $r$ is the rate of change, and $t$ is the elapsed time. This equation is solved for $r$ such that:

$$r = ln(N/N_0)/t \quad \text{(eqn. S11)}$$

We then calculated population growth rates using values for the first and last observed values for each process. To verify results, we integrated values for $r$, $t$, and $N_0$ to eqn. S7 and compared results to observed values for $N$ (Table S4).

#### 6. Catch rate analyses

We compared catch rates, measured as mean annual CPUE, in commercial fishing (1952-1982; LEK data) and scientific monitoring (1995-2018; Monitoring data). We used a Mann-Whitney U test, as neither dataset was normally distributed (Table S5).

#### 7. Size distribution analyses

We analyzed size distribution by evaluating Curved Carapace Length (CCL, cm), and by grouping juveniles and adults using the mean CCL of nesting females at Colola (82.0 cm) (Alvarado Díaz & Figueroa cited in Seminoff et al. 2015). We divided monitoring data into two groups according to data availability: Period 1 (1995-2005) and Period 2 (2009-2018). We obtained descriptive statistics of CCL in both periods (Table S6). Given the non-normal distribution of CCL values in Period 2, we used a Mann-Whitney U test ($\alpha = 0.05$) to compare size composition between periods, and found significant differences (Table S7).

Early-Capistrán et al. (2021) Pre-print

### Figures

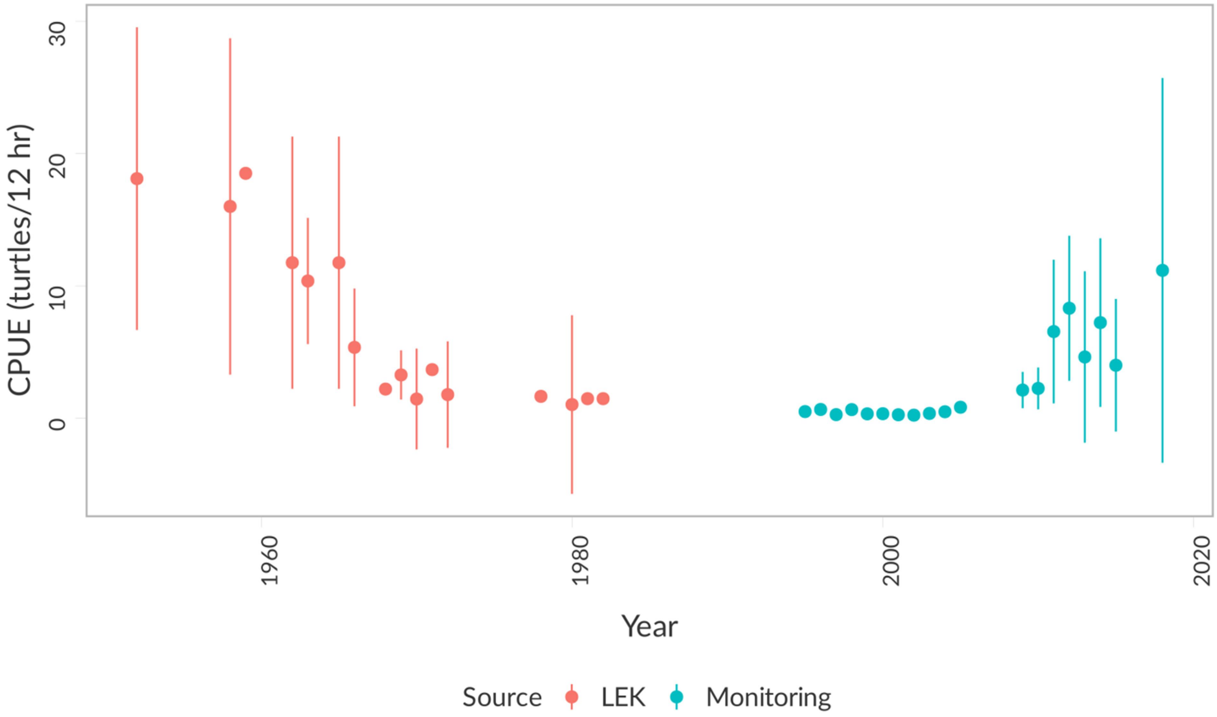

**Figure S1: All CPUE values from LEK and Monitoring datasets.**

All CPUE values from LEK (1952-1982) and Monitoring (1995-2018) datasets, with means (points) and 95% Confidence Intervals. Note that some years had only one observation. Mean annual CPUE values were used for all analyses. See also Tables S1, S2.

Early-Capistrán et al.

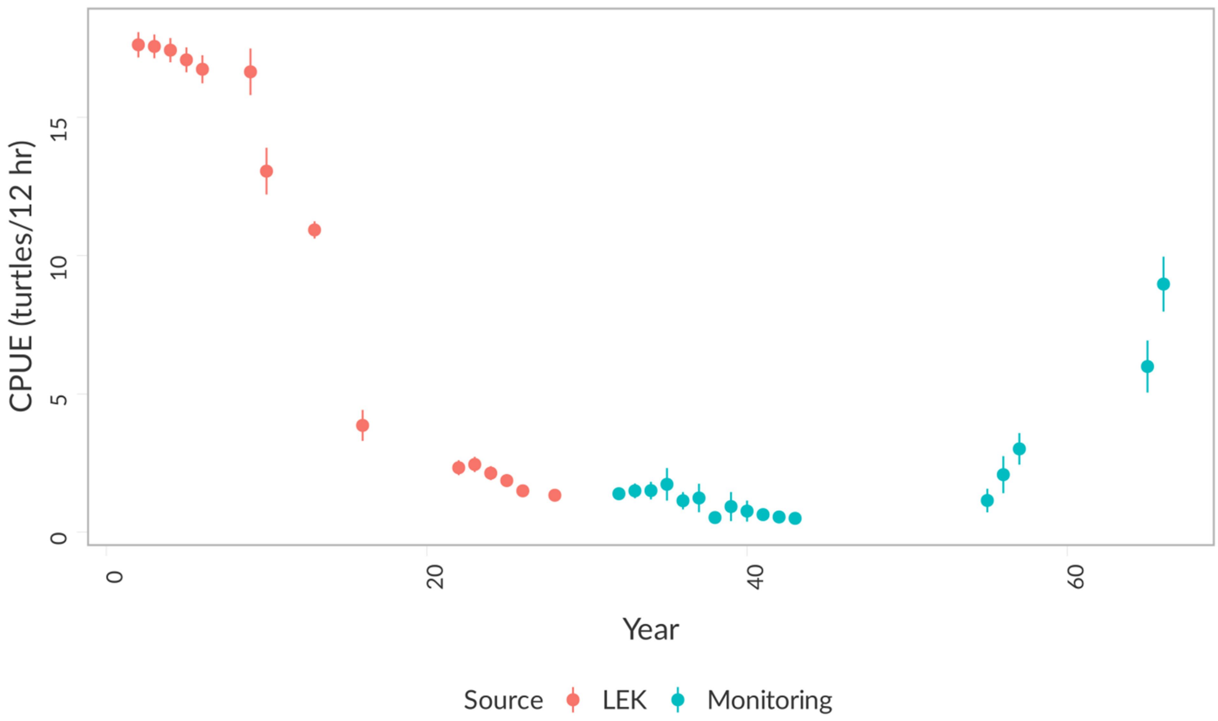

**Figure S2: Mean multiply imputed CPUE values**

Mean multiply imputed CPUE values (points) with 95% Confidence Intervals. Note that narrow confidence intervals are due to the large number of imputed data points per year, sampled from subsets of observed data. See also Figure S3.

Early-Capistrán et al. ©

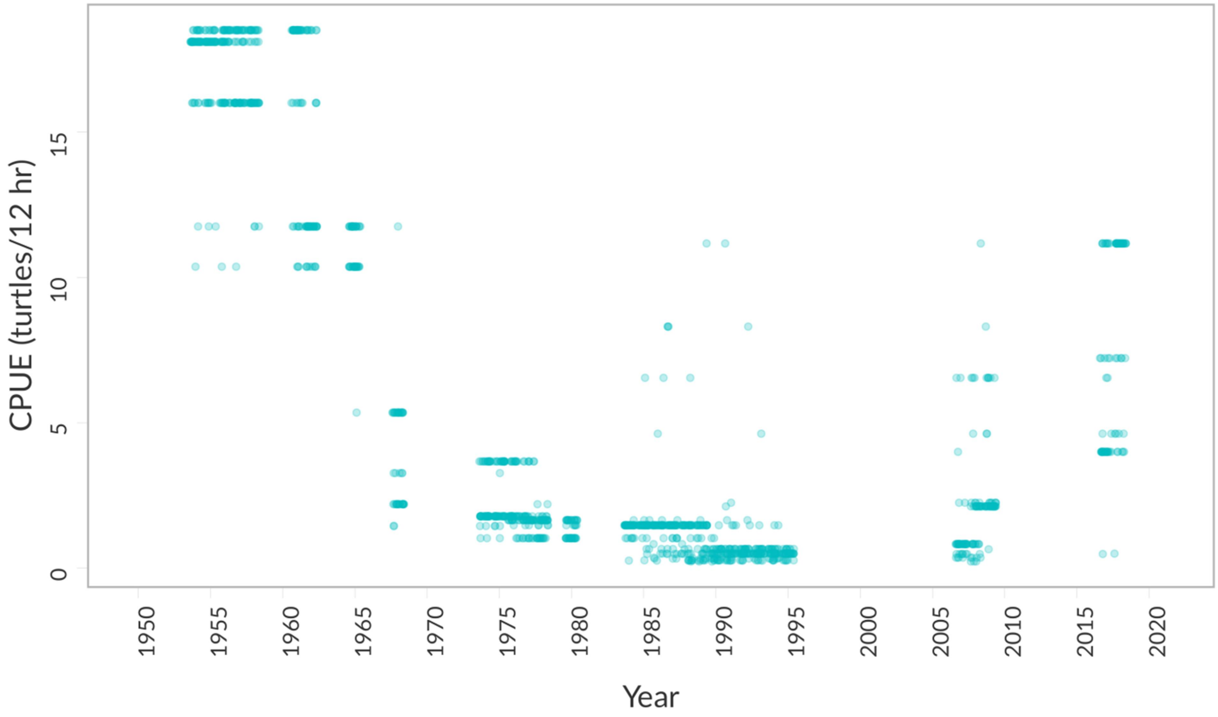

**Figure S3: Scatterplot of all imputed values**

All values imputed for missing data from LEK and Monitoring datasets (N = 1434).

Early-Capistrán et al.

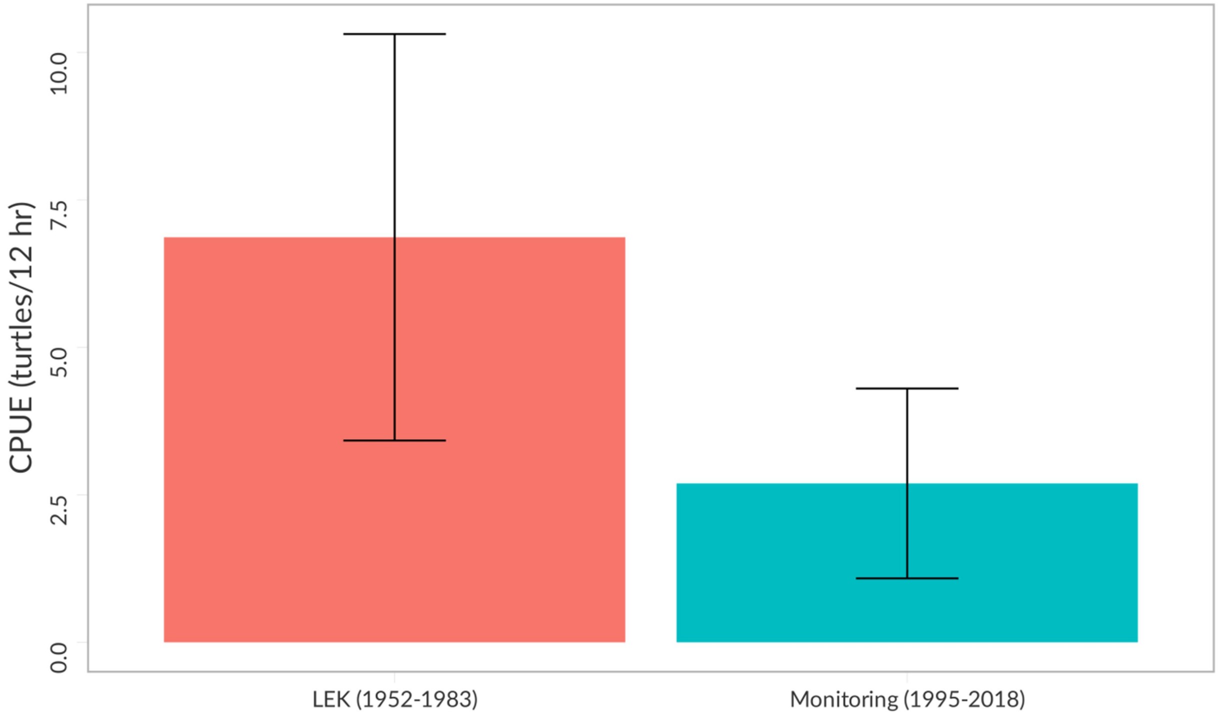

**Figure S4: Mean annual CPUE values grouped by dataset**

Mean annual catch-per-unit-effort (CPUE) values grouped by dataset (LEK and Monitoring), with error bars showing 95% Confidence Intervals. See also Table S5.

Early-Capistrán et al.

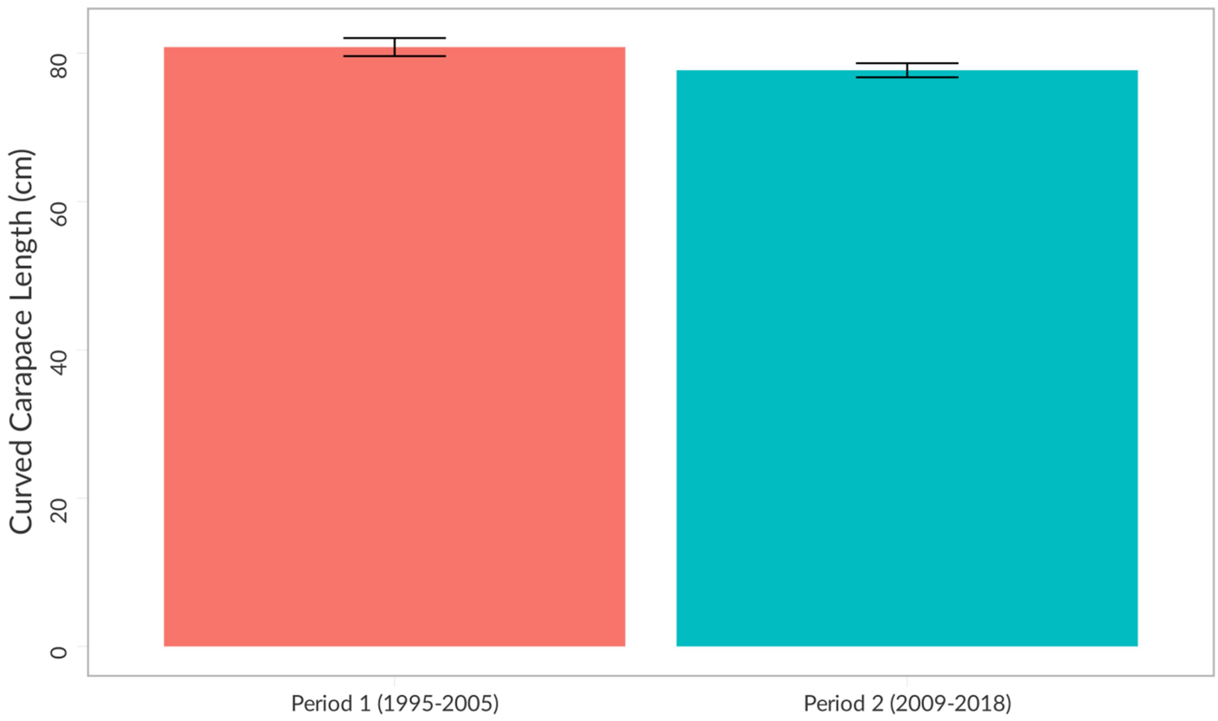

**Figure S5: Curved Carapace length grouped by period**

Curved Carapace Length (CCL, cm) in Monitoring data, grouped by period, with error bars showing 95% Confidence Intervals. See also Table S7.

Early-Capistrán et al.

### Tables

**Table S1**: Descriptive statistics of all catch-per-unit-effort (CPUE) values

| Dataset | N | Mean | Median | Standard deviation | Minimum | Maximum |
|---|---|---|---|---|---|---|
| LEK data (1952-1982) | 32 | 6.68 | 3.91 | 6.04 | 0.5 | 19 |
| Monitoring data (1995-2018) | 148 | 3.21 | 1 | 6.45 | 0 | 36 |

**Table S1: Descriptive statistics of all CPUE values**

Early-Capistrán et al. (2021) Preprint

**Table S2**: Descriptive statistics of mean annual catch-per-unit-effort (CPUE) values

| Dataset | N | Mean | Median | Standard deviation | Minimum | Maximum |
|---|---|---|---|---|---|---|
| LEK data (1952-1982) | 16 | 6.86 | 3.47 | 6.46 | 1.03 | 18.5 |
| Monitoring data (1995-2018) | 19 | 2.69 | 0.66 | 3.34 | 0.23 | 11.17 |

#### Table S2: Descriptive statistics of mean annual CPUE values

Early-Capistrán et al. (2021) Preprint

**Table S3:** Results of nonlinear regression for mean annual catch-per-unit-effort. Italics indicate significant results at $\alpha = 0.05$.

| Parameter | Estimate | Std. Error | 95% C.I. | *t*-value | P-value | $R^2$ |
|---|---|---|---|---|---|---|
| LEK data: commercial fishing (1952-1982), nonlinear regression; Model: $y \sim \alpha^{(\beta \cdot x)}$; $df = 14$; $e \sim N(0, \sigma^2)$ | | | | | | |
| α | 24.112 | 3.124 | [17.413 -30.812] | 7.719 | *2.07e-06* | 0.798 |
| β | -0.0829 | 0.0130 | [-0.111 to -0.0551] | 6.382 | *1.71e-05* | |
| Monitoring data: conservation (1978-2018*), nonlinear regression; Model: $y \sim \alpha^{(\beta \cdot x)}$; $df = 21$; $e \sim N(0, \sigma^2)$ | | | | | | |
| α | 5.40e-04 | 8.22e-04 | [-0.00117 - 0.00225] | 0.657 | 0.518 | 0.780 |
| β | 0.148 | 0.0238 | [0.0985 - 0.198] | 6.212 | *3.67e-06* | |

* LEK values from 1978-1982 were appended to interpolate for the temporal gap between the end of commercial fishing (1983) and the start of in-water monitoring (1995).

#### Table S3: Results of non-linear regression for mean CPUE

Results of nonlinear regression for mean annual catch-per-unit-effort (CPUE). Italics indicate significant results at $\alpha = 0.05$.

Early-Capistrán et al. (2021) Pre-p

**Table S4:** Annual population growth rates during commercial fishing and conservation

| Phase | Start year | End year | Annual growth rate | Predicted value | Observed value |
|---|---|---|---|---|---|
| Commercial Fishing<br><br>(LEK data) | 1952 | 1982 | -8.4% | 1.456 | 1.475 |
| Conservation<br><br>(LEK and Monitoring data) | 1978* | 2018 | 4.8% | 11.075 | 11.167 |

* As in MICE analyses, LEK values from 1978-1982 were appended to interpolate for the temporal gap between the end of commercial fishing (1983) and the start of in-water monitoring (1995). See also Figure 1.

#### Table S4: Annual population growth rates

Annual population growth rates during Commercial Fishing and Conservation phases. Growth rates were calculated using mean annual catch-per-unit-effort (CPUE) values.

Early-Capistrán et al. (2021) Pre-p

**Table S5**: Results of Mann-Whitney U test for mean annual catch-per-unit-effort (CPUE) values in LEK and monitoring datase. Italics indicate significant result (α = 0.05).

| Variable | Period | N | U | p | 95% C.I. |
|---|---|---|---|---|---|
| CPUE (turtles/night) | LEK (1952-1982) | 16 | 232 | *0.008* | 0.81 – 7.75 |
| | Monitoring (1995-2018) | 19 | | | |

#### Table S5: Mann-Whitney U test for CPUE values

Results of Mann-Whitney U test for mean annual catch-per-unit-effort (CPUE) values in LEK and Monitoring datasets. Italics indicate significant result (α = 0.05).

Early-Capistrán et al. (2021) Pre-print

**Table S6:** Descriptive statistics of size distribution (Curved Carapace Length, cm) in monitoring data

| Period | N | Mean | Median | Standard deviation | Minimum | Maximum |
|---|---|---|---|---|---|---|
| Period 1 (1995-2005) | 289 | 80.85 | 80.8 | 10.51 | 51.1 | 104.5 |
| Period 2 (2009-2018) | 414 | 77.72 | 75.5 | 9.83 | 54.5 | 109 |

#### Table S6: Descriptive statistics for Curved Carapace Length

Descriptive statistics of size distribution (Curved Carapace Length, cm) in monitoring data.

Early-Capistrán et al. (2021) Pre-p

**Table S7**: Results of Mann-Whitney U test for Curved Carapace Length (CCL) in monitoring data. Italics indicate significant result (α = 0.05).

| Variable | Period | N | U | p | 95% C.I. |
|---|---|---|---|---|---|
| Straight Carapace Length (cm) | Period 1 (1995-2005) | 282 | 71406 | *1.228e-05* | 1.99 – 5.20 |
| | Period 2 (2009-2018) | 414 | | | |

#### Table S7: Mann-Whitney U test for Curved Carapace Length.

Results of Mann-Whitney U test for Curved Carapace Length (CCL) in monitoring data. Italics indicate significant result (α = 0.05).

Early-Capistrán et al. (2021) Pre-print

#### References

Barnard, J. & Rubin, D.B. (1999). Small-Sample Degrees of Freedom with Multiple Imputation. *Biometrika*, 86, 948–955.

Bodner, T.E. (2008). What Improves with Increased Missing Data Imputations? *Structural Equation Modeling: A Multidisciplinary Journal*, 15, 651–675.

van Buuren, S. & Groothuis-Oudshoorn, K. (2011). MICE : Multivariate Imputation by Chained Equations in R. *Journal of Statistical Software*, 45.

Dong, Y. & Peng, C.-Y.J. (2013). Principled missing data methods for researchers. *SpringerPlus*, 2, 222.

Early-Capistrán, M.-M., Solana-Arellano, E., Abreu-Grobois, F.A., Narchi, N.E., Garibay-Melo, G., Seminoff, J.A., Koch, V. & Saenz-Arroyo, A. (2020). Quantifying local ecological knowledge to model historical abundance of long-lived, heavily-exploited fauna. *PeerJ*, 8, e9494.

Jia, F. (2016). *Methods for Handling Missing Non-Normal Data in Structural Equation Modeling* (PhD).

Nguyen, C.D., Carlin, J.B. & Lee, K.J. (2017). Model checking in multiple imputation: an overview and case study. *Emerging Themes in Epidemiology*, 14.

Rubin, D.B. (1987). *Multiple imputation for nonresponse in surveys*. Wiley series in probability and mathematical statistics. Wiley, New York.

Santacruz López, E. (2012). *Monitoreo estandarizado de la población de tortugas marinas, en la Reserva de la Biosfera Bahía de Los Ángeles, Baja California, México* (Maestría en Ciencias).

Early-Capistrán et al. (2021) Pre-print

Seminoff, J.A., Allen, C.D., Balazs, G., Dutton, P.H., Eguchi, T., Haas, H.L., Hargrove, S., Jensen, M., Klemm, D.L., Lauritsen, A.M., MacPherson, S.L., Opay, P., Possardt, E.E., Pultz, S., Seney, E., Van Houtan, K.S. & Waples, R.S. (2015). *Status Review of the Green Turtle (Chelonia mydas) Under the Endangered Species Act* (NOAA Technical Memorandum No. NOAA-TM-NMFS-SWFSC-539). NOAA, San Diego, USA.

Seminoff, J.A., Jones, T.T., Resendiz, A., Nichols, W.J. & Chaloupka, M.Y. (2003). Monitoring green turtles (*Chelonia mydas*) at a coastal foraging area in Baja California, Mexico: multiple indices to describe population status. *Journal of the Marine Biological Association of the UK*, 83, 1355–1362.

Seminoff, J.A., Reséndiz-Hidalgo, A., Jiménez de Reséndiz, B., Nichols, W.J. & Todd-Jones, T. (2008). Tortugas marinas. In: *Bahía de los Ángeles: recursos naturales y comunidad: línea base 2007* (eds. Danemann, G. & Ezcurra, E.). Secretaría de Medio Ambiente y Recursos Naturales ; San Diego Natural History Museum, Tlalpan, México D.F.; San Diego, Calif., pp. 457–494.

van Buuren. (2018). *Flexible Imputation of Missing Data*. Chapman & Hall/CRC Interdisciplinary Statistics Series. 2nd edn. CRC Press, New York.

Early-Capistrán et al. (2021) Pre-print
